# A Test of Memory: The Fish, The Mouse, The Fly And The Human

**DOI:** 10.1101/2020.02.15.950816

**Authors:** Madeleine Cleal, Barbara D Fontana, Daniel C Ranson, Sebastian D McBride, Jerome D Swinny, Edward S Redhead, Matthew O Parker

## Abstract

Simple mazes have provided numerous tasks for assessing working memory. The discrete nature of choices in the T-maze has provided a robust protocol with sensitivity to cognitive deficits, whilst the continuous Y-maze reduces manual handling and pre-trial training. We have combined these attributes to develop a new behavioural task for assessing working memory, the Free-movement pattern (FMP) Y-maze. Using sequentially recorded left and right turns we demonstrate that zebrafish and mice use a single dominant strategy predominantly consisting of alternations between left and right choices trial-to-trial. We further tested this protocol with *Drosophila* and discovered an alternative invertebrate search strategy. Finally, a virtual human FMP Y-maze confirmed a common strategy among all tested vertebrate species, validating the translational power of the task for human research. The FMP Y-maze combines robust investigation of working memory and high translational power, generating a simple task with far-reaching impact.

## Introduction

Mazes have provided invaluable insight into the many ways in which short-term and working memory are utilised by animals^1^. However, conventional set ups can often require extensive training, high levels of animal handling and, in reward-based trials, food or water deprivation for prolonged periods. Each of these factors can result in potential confounders, leading to high levels of between-subject variability^2^. Additionally, not all tests can easily be adapted between different species, and with the rise of zebrafish as a model organism for neuropsychiatric research, effective assays need greater flexibility to enable use with a range of organisms^3^. Exploiting the neophilic nature of rodents, an unbiased T-maze is one of the simplest mazes used to assess working memory^4^. However, discrete versions of the task typically require a substantial degree of animal handling, which, although only mildly stressful for land animals, can potentially cause distress or injury for aquatic organisms that require netting/chasing or air exposure to return them to the beginning of the task^2^. This level of intervention could reasonably be presumed to affect an animal’s search strategy in subsequent trials.

Similar to the T-maze, the Y-maze, has demonstrated great promise for aquatic organisms due to the continuous, non-reinforced nature, which revokes the need for unnecessary handling or pre-trial training, making this one of the quickest of the maze tests ^5,6^. However, there appear to be limitations, in a review of different memory tasks ability to detect differences between controls and the Tg2576 Alzheimer’s mice, the continuous Y-maze was one of the worst performers, with only 53% of studies analysed able to detect differences between controls and Tg2576 mice ^7^. Despite the short run time making this task popular for assessing working memory, (Hughes, 2004) highlights practical concerns resulting in an unfavourable view of the continuous Y-maze. Ultimately, both (Hughes, 2004 and Stewart, Caccui and Lever, 2011) imply a lack of suitability as a stand-alone memory test, thus, requiring it to be run as part of a battery of behavioural tasks ^7,8^.

To address these issues we have developed The Free-movement pattern (FMP) Y-maze, which has been designed to combine the proficiency of the T-maze with the timesaving, non-reinforcing nature of the continuous Y-maze. Based on the protocol of the two-choice guessing task ^9,10^, we systematically report a time series of left and right turns made by a subject whilst navigating the maze. To the contrary of common Y-maze methods, this protocol was not aimed at assessing response to novelty, but to identify if exploration of a restricted area was conducted at random or using distinct search patterns. Based on the work by (Frith and Done, 1983) turn choices were divided into quadruplets (sequences of four turns known as a tetragrams) in order to exploit the information theory measure to identify randomness in large sequences of data^10^. We furthered this analysis by using autocorrelation function to examine if a tetragram sequence at position *i* in a time series influenced subsequent tetragram choices. To validate this task as test of memory, we pre-treated zebrafish with antagonists known to inhibit memory formation. If search strategies employed in the FMP Y-maze are dependent on memory processing and formation, we hypothesis that blocking these pathways would disrupt normal searching behaviour in favour of a less structured strategy. Once a subject has been placed into the maze the task is run for a set time, under semi-dark conditions, limiting the effects of egocentric cues and experimenter interference^11^. Here, we present a system that demonstrates sensitivity to behavioural changes resulting from the disruption of memory forming pathways and overcomes limitations of species specificity, excessive handling, reward biases and pre-trial training that hinder many other maze-based memory tests. In addition, this task has easily been adapted to a virtual FMP Y-maze for humans for the ultimate translatability from model organism to patient. This task enables previously unnoticed choice-behaviours to be identified in both vertebrates and invertebrates, with distinct overlap in the search strategies of vertebrate model organisms and humans. Thus, the FMP Y-maze demonstrates its suitability as a stand-alone test of memory for a vast range of organisms.

## Results

### Implementation of the FMP Y-maze

The FMP Y-maze is a simple design that has been incorporated into an automated tracking system. The Zantiks behavioural unit provides a partially enclosed environment which both shields experimenters from view, limiting distraction whilst animals are in the maze, and maintains dimly lit conditions, hampering dependence on external cues for navigation **(Fig. 1A-B)**. This set up simultaneously allows the free movement of animals and the automated logging of arm entries and exits uninterrupted for the duration of the trial. The mazes were designed so each model (fish, mouse or fly) had equal arm length (relative to body size) and angle, to avoid unintended arm biases during exploration **(Fig. 1C_Supplementary_Fig. 1)**. Tetragram analysis was based on 16 overlapping sequences of left and/or right turns, each representing a unique sequence of arm entries in the FMP Y-Maze **(Fig. 1D_Supplementary_Table. 1, 1.3_Tetragram analysis)**. Heat maps and line traces were recorded for a random selection of individuals from each species to identify any arm bias when searching the maze **(Fig. 1E-G, Fig. 3B,C,E,F_Supplementary Fig. 2)**. Hot spots in select arms likely represented ‘rest’ periods between searching bouts. Line traces of movement trajectories show equal use of all arms and no significant biases were observed when analysing the proportion of entries in each arm (one-way ANOVA; F(2, 48) = 2.877; P=0.0661; n=17).

**Figure 1.**
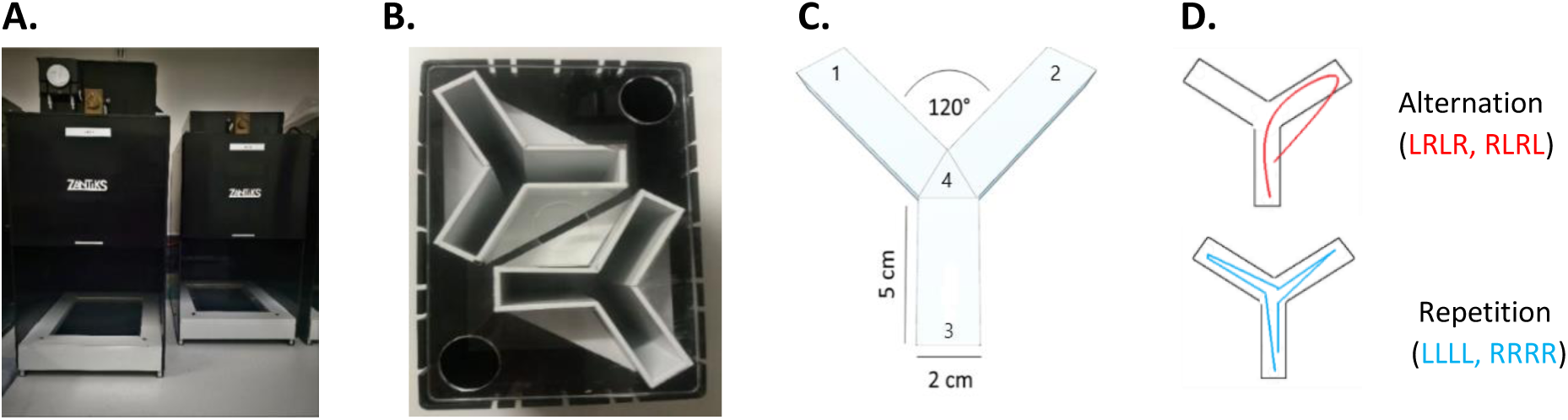
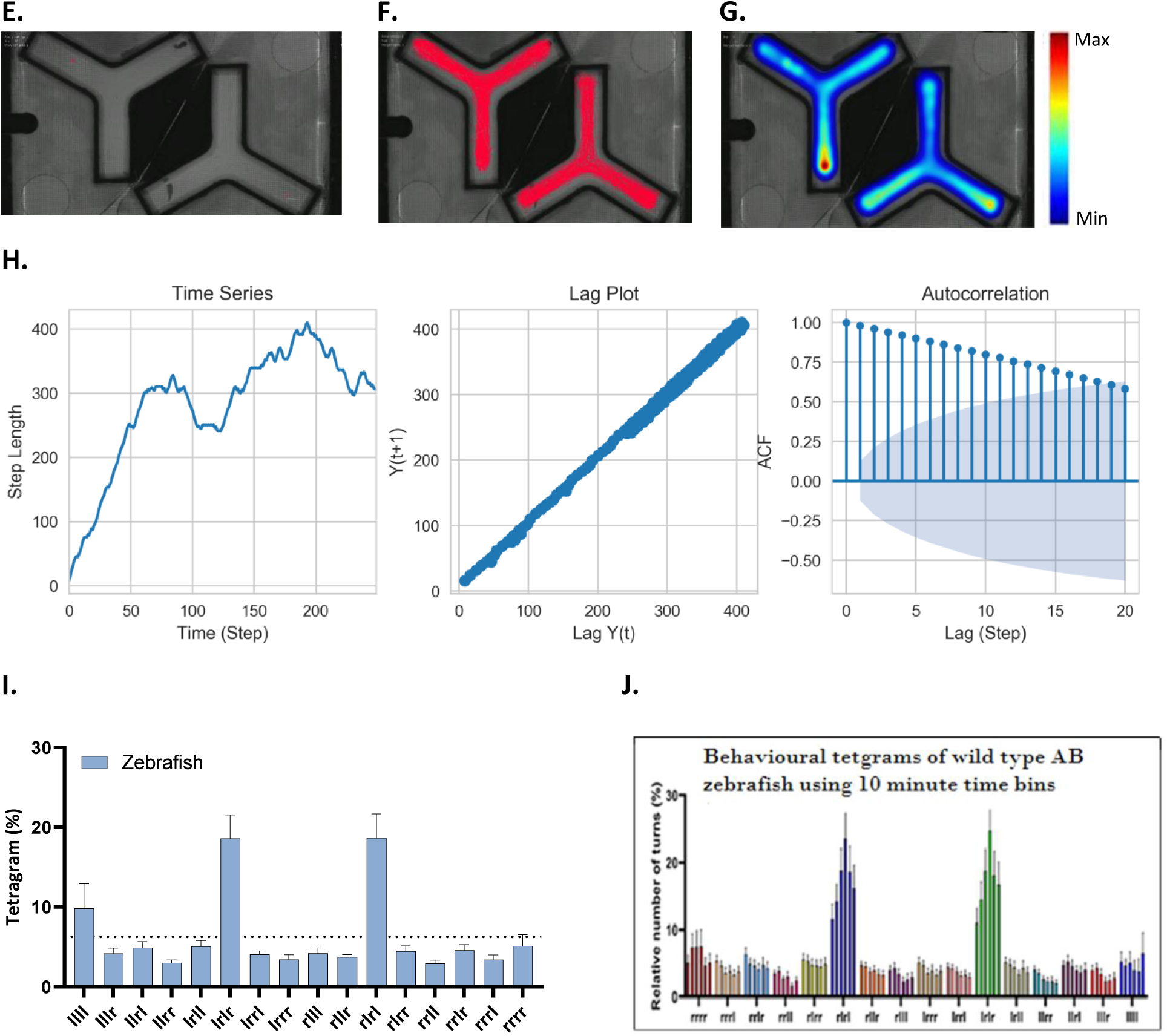
Aquatic FMP Y-maze for zebrafish. (**A**) Zantiks behavioural unit for automated animal tracking. (**B**) Top view of two FMP Y-mazes for zebrafish inserted into a black water-tight tank, L50:W20:H140mm, filled with 3L of aquarium water. A mesh lid was used to cover the top of the tank to prevent fish from jumping out during the trial without interfering with the tracking software. (**C**) FMP Y-maze diagram depicting maze dimensions and zones used for automated logging of arm entries and exits. (**D**) Schematic illustrating movement patterns, in the form of line drawings of arm entries and exits, used for two specific tetragram sequences, alternations (LRLR, RLRL) and repetitions (LLLL, RRRR). (**E**) In trial image of zebrafish in the FMP Y-maze (n=2). (**F**) Line trace of 1 h of locomotor activity (n=2). (**G**) Heat map analysis of animal tracking following 1 h of exploration time (n=2). Red represents increased time spent and blue represents minimal time spent during trial. (**H**) Time series analysis of movement patterns of individual zf11 (n=1), showing from left to right, time series plot of the cumulative sum of step lengths for n=250 time points. Lag plot of data at lag-0 (ω(*k*)) and lag-1 (ω(*k*+1) demonstrating a positive linear correlation. Autocorrelation function plot showing the first 20 lags of 250 lag plot. Plot shows slow decay towards zero, with 18 lag points outside of the 95% C.I., depicted by the blue cone. Autocorrelation between data points is indicative dependency between successive turn choices, demonstrating memory of previous choices. **(Supplementary_Table.2).** Error bars represent ± SEM.

**Figure 2.**
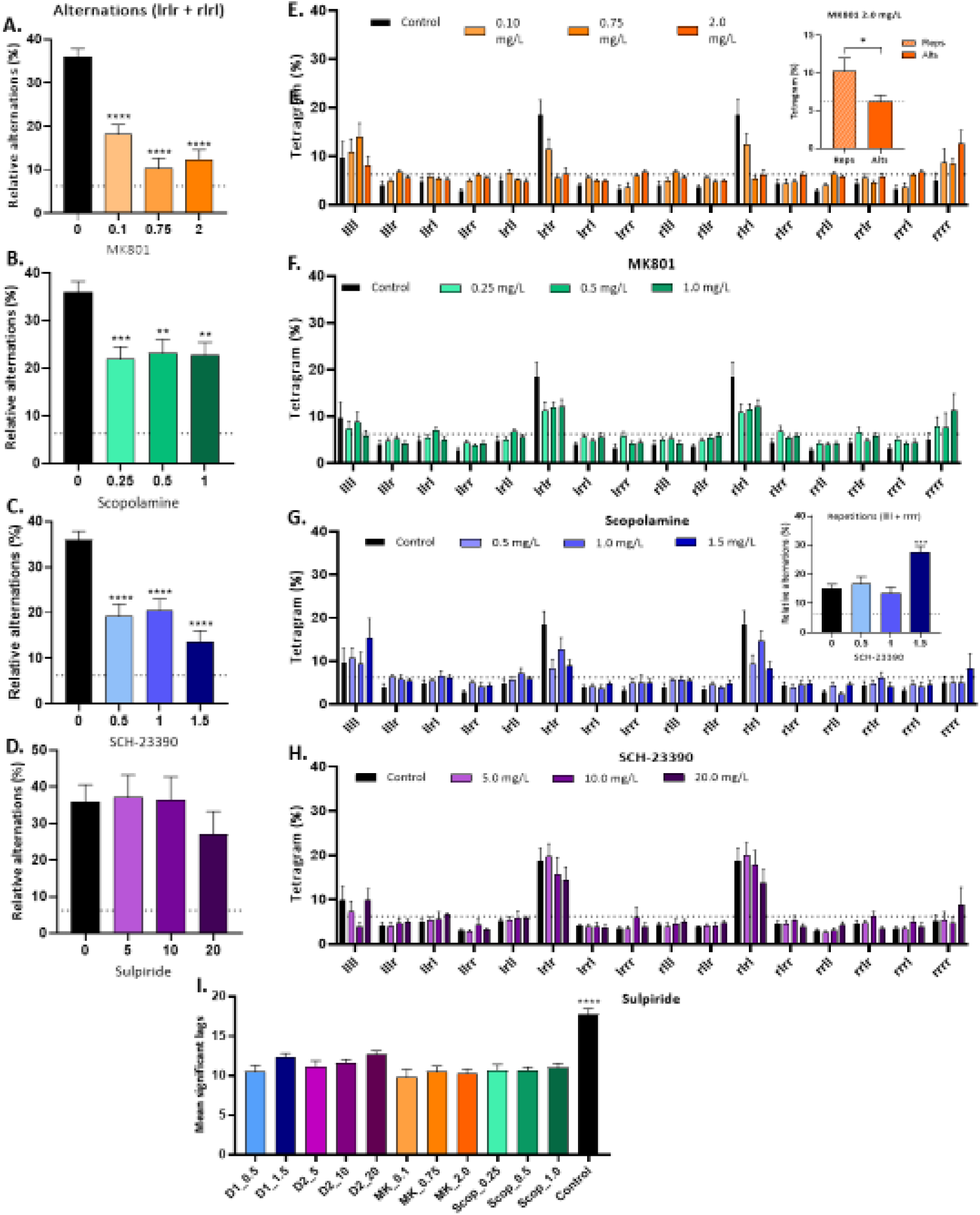
Effect of pharmacological treatment on movement patterns of zebrafish in the FMP Y-maze. (**A-D**) total percentage use of combined alternation strategies (LRLR + RLRL) during a 1 h exploration trial in the FMP Y-maze. Each drug and dose were compared to the mean (µ=35.989) of the control group (n=18). All doses of MK801 (0.1, 0.75 and 2.0 mg/L, n=13 per group) showed significant decrease of alternations (P<0.0001) (A). Scopolamine (0.25, 0.5 and 1.0 mg/L, n=13 per group) showed the largest decrease in alternations at the lowest dose (0.25 mg/L) (P<0.0001) and a reduced decrease at the mid and high doses (P<0.001) (B). SCH-23390 showed similar decrease in alternations to MK801 (0.5, 1.0, 1.5 mg/L, n=12 per group) (P<0.0001) (C). Sulpiride showed no significant effect on alternations (P=0.590) (D). (**E-H**) Percentage use of each of the 16 tetragram sequences over the duration of a 1 h trial for low, mid and high doses for each drug compared to the control. (E) insert, the highest dose of MK801 (2.0 mg/L) shows alternations were used less than repetitions throughout the trial (P=0.0364). (G) insert, high dose of SCH-23390 caused a significant increase in the use of the repetition strategy (LLLL + RRRR) compared to control fish (P<0.0001). (**I**) ACF plots for each individual fish were computed in MATLAB **(Supplementary_Time series analysis, Figure 7-10, Table.3)**, mean number of lag points with significant autocorrelation were compared to mean of controls. All drug treated animals showed a significant increase in decay rate (P<0.0001) showing a reduced influence of past choices on lag(k). Error bars represent ± SEM.

**Figure 3.**
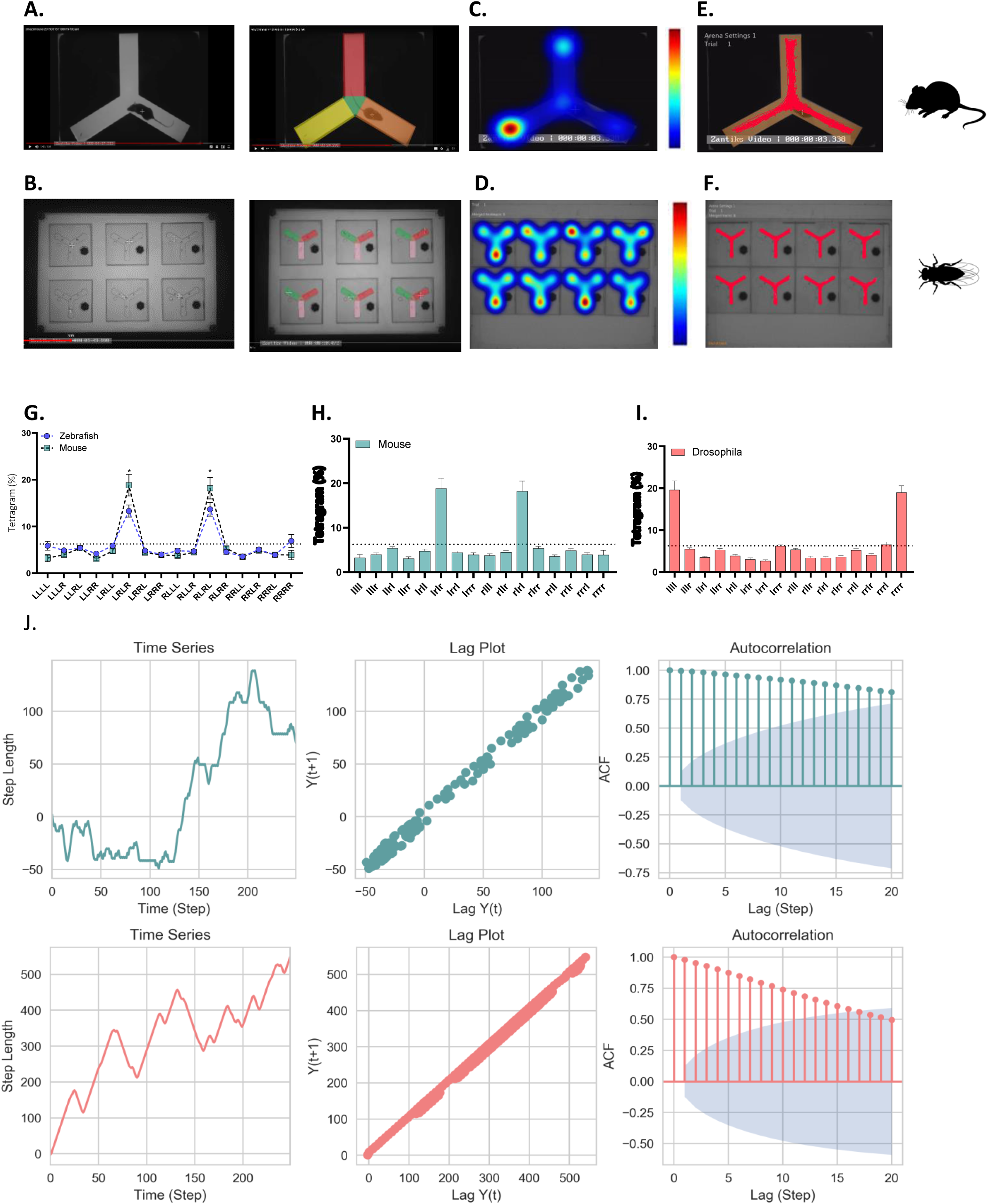
FMP Y-maze exploration strategies of mice and *Drosophila*. (**A-B**) In maze view of a single mouse in the rodent FMP Y-maze (top, n=1) and 6 individual fly mazes each containing a single *Drosophila* (bottom, n=6) with and without zone markings. (**C-D**) Heat map analysis of animal tracking following 1 h of exploration time (n=1 mouse, n=8 *Drosophila*). Red represents increased time spent and blue represents minimal time spent during trial, measured using Ethovision software of video tracking. (**E-F**) Line trace from 1 h exploration trial in the rodent and fly mazes, respectively. (**G**) Overlay of tetragram usage throughout a 1 h trial of mice compared to zebrafish exploration (n=15 mice, n=18 zebrafish) demonstrating almost identical strategies, the only deviation was between the percentage use of alternations which was greater in rodents (P=0.0272). (**H-I**) Frequency distribution of tetragram use over a 1 h trial (n=15 mice, n=30 *Drosophila*) with mice showing preferential use of alternations and *Drosophila* dominantly using repetitions. (**J**) Time series analysis of an individual mouse (top) and *Drosophila* (bottom) showing time series plot of step length (n=250 steps), lag plot showing positive correlation for both organisms, ACF plot of the first 20 lags, both demonstrating over 15 lags of significant autocorrelation. Error bars represent ± SEM.

### Validation of the FMP Y-maze in an aquatic environment

We developed a system to determine the exploration strategy of zebrafish in an unknown environment. Without prior training or habituation, fish were permitted to freely explore the arena for 1 h with continuous recording of arm entries and exits for the duration of the trial. The absence of reinforcement means that fish do not require periods of pre-trial food deprivation and can therefore, be taken directly from home tank to test tank and back to home tank, keeping handling and stress to a minimum in accordance with the 3Rs. Our primary aim was to identify if the FMP Y-maze could be used as a test of memory. Data from the task was output as a discrete time series^12,13^, from which it is possible to mathematically model the randomness of serial observations ^14^. First, we applied the two-choice guessing task system, utilised by (Frith and Done, 1983), of tetragram analysis to identify any discernible patterns within the movements that departed from a random process^9,10^. The sum of each of 16 overlapping tetragrams of left and/or right turns (e.g. left, left, left, left [L,L,L,L] or right, right, left, left [R,R,L,L]) were averaged for the group, revealing a strategy that showed significant use of one particular type of tetragram sequence containing alternating left and right turns (LRLR, RLRL), referred to from here as alternations **(Fig. 1D)** (one-way ANOVA; F(15, 272) = 17.31; P<0.0001; n=21). Although similar to the alternating pattern from the T-maze, in the FMP Y-maze alternations were not used exclusively (which might be consistent with stereotypic behaviour), but instead were distributed regularly throughout the trial **(Fig. 1J-K)**. Alternations were used as a search strategy ∼38% of the time, regularly dispersed with other combinations of the remaining 14 tetragrams. The regular occurrence of a specific type of tetragram, the alternation, indicates a complex level of behaviour in which the preceding trigram sequences LRL or RLR, are predictors that the following turn choice will be a R or L turn respectively, demonstrating strong intrasequence dependencies. Thus, despite the overall probability of turning L or R being equally likely, the use of tetragram analysis has revealed the presence of a repeating pattern within the data, resulting in a deviation from complete randomness. Although tetragram analysis can be used to identify preferential turn choices and dependency of a choice based on the three preceding turns, it cannot be used to determine the persistence of that dependency. Put simply: for a turn choice at position *i*, to what extent are subsequent turns influenced? Using the lag-1 autocorrelation function (ACF) it is possible to determine the relationship between successive tetragram sequences and identify if dependency lasts beyond the tetragram set.^15^ ACFs that rapidly decay, fluctuating around zero, would be indicative of a completely random, or memoryless process^16^, i.e. a Markovian process^17^. However, as we have already demonstrated strong intrasequence dependency of particular tetragrams, we know that the sequence of turn choice are not random. However, there is no indication of whether dependency of prior turns lasts only for three previous turn choices (a tetragram sequence) or perpetuate for longer periods (turn choices equivalent to multiple tetragram sequences). Our evidence strongly suggests that movement patterns were the result of a specific strategy, relying on memory of past turn choices. We therefore hypothesised that subsequent steps (each step representing a tetragram) would demonstrate significant autocorrelation, which would be suggestive of a time series with *memory* of previous events, which exert influence on choice-behaviour for a large number of steps. We found that time series plots for individual zebrafish showed either left or right bias, but the ACF of the cumulative sum of steps showed prolonged autocorrelation, which decayed slowly to zero **(Fig. 1H-I_ Supplementary_Time series analysis Fig. 3)**. These ‘long-range correlations’ between turn choices reflect a long-lasting effect of previous behaviour on subsequent choice-behaviour. In sum, our data suggest that the generation of the behavioural sequences of turns by wild type adult zebrafish in the FMP Y-maze are characterised by long-range and significant non-random relationships between steps across a large range of responses **(Supplementary_Video.1)** and that this sequence of choice behaviour is based on the memory of previous actions.

### Changes in exploration strategy due to pharmacological blockade of memory pathways

To substantiate the use of memory to strategically navigate the FMP Y-maze we pharmacologically targeted discrete pathways of the memory processing system. Blocking pathways involved in encoding or consolidating memories would be expected to alter choice selection in the FMP Y-maze. To this end, we pre-treated zebrafish with a high, mid and low concentration of four antagonists, inhibiting key receptors in the memory process. MK-801, a non-competitive NMDA receptor (NMDA-r) antagonist known to interfere with memory formation by inhibiting long-term potentiation (LTP)^18,19^, caused a significant decrease in the use of alternations compared to control fish (GLMM, F(3, 318) = 34.221; P < 0.0001, 0.1 mg/L n=13, 0.75 mg/L n=13 and 2.0 mg/L n=13, control n= 18). If each tetragram was used equally throughout the trial, use would be expected at ∼6.25%, equivalent to random selection of each tetragram sequence. Wild type zebrafish exhibit the use of alternations at ∼38% showing significant non-randomness. However, treatment with MK801 reduces the use of alternations to less than 6%, effectively blocking use of alternations as a strategy. At the highest dose (2.0 mg/L) a role reversal was evident as repetitions were used to a greater extent than alternations (T-test; t(2.155, 46), P=0.0364, n=24) **(Fig. 2A-B)**. Scopolamine, a non-specific muscarinic receptor (M-r) antagonist, similarly to MK-801 behaves by reducing LTP in the hippocampus^20^, also decreased the use of alternations to navigate the maze, but to a lesser extent than MK-801 (GLMM, F(3, 316) = 8.025; P < 0.0001, 0.25mg/L, n=13; P<0.001, 0.5, n=13 and 1.0 mg/L, n=13) **(Fig. 2C-D)**. The dopaminergic system is also critical to learning and memory formation^21^. Divided into two sub-groups, D1-like (D1 and D5 receptors) and D2-like (D2, D3 and D4 receptors) receptors are crucial to working memory and cognition-D1-like-and motivation and reward learning-D2-like^22^. Treatment with SCH-23390, a D1-like receptor antagonist, caused two major changes in search strategy. At all concentrations, there was a decrease in the use of alternations, similarly to that caused by MK-801. Additionally, the highest concentration caused an increase in the use of an another tetragram coupling consisting either entirely of left or right turns (LLLL, RRRR), described as pure repetition (GLMM test, F(3, 311) =19.692; P< 0.0001, 0.5 mg/L, n=12; 1.0 mg/L, n=12; 1.5 mg/L, n=12. GLMM test, F (3, 312) = 8.954; P<0.001, 1.5 mg/L, n=12). There was no significant difference between the use of alternations and any other tetragram configuration at 0.5 and 1.5 mg/L **(Fig. 2E-F)**. No such effect was evident with the D2-like antagonist, sulpiride, which was associated with no change in strategy compared to control fish (GLMM test, n=33, P=0.622) **(Fig. 2G-H)**. ACF plots of each concentration of drug resulted in a decrease in the number of significantly correlated lags compared to control fish (one-way ANOVA; F(11, 127) = 13.94; P<0.0001) **(Fig. 2I_Supplementary_Figure7-10)**. Thus, memory impaired zebrafish resulted in shorter-range correlations, limiting the number of steps influenced by choice behaviours showing a reduction in information processing capabilities compared to controls.

### Validating the FMP Y-maze with other model organisms-the mouse and the fly

Having demonstrated the suitability of the FMP Y-maze for assessing fish, we tested the generalizability of the test to other widely used laboratory species (mouse and *Drosophila*) in order to characterise exploration strategies used to navigate the FMP Y-maze for 1 h, following an identical protocol to zebrafish **(Fig. 3A, D_Supplementary_Figure 1A,C)**. We observed two key findings. Mice navigated the FMP Y-maze using an almost identical strategy to zebrafish, showing dominant use of alternations throughout the task **(Fig. 3G-H)**. *Drosophila* differed from mice and zebrafish, however, employing an exploration strategy that was not reliant on alternations but on pure repetitions, with these accounting for ∼38% of their total search strategy **(Fig. 3I)**. This alternative navigational pattern could be influenced by *Drosophila*’s natural tendency to explore using wall-following behaviour^23^. Like mice and zebrafish, *Drosophila* displayed their dominant strategy at evenly distributed times throughout the task, regularly interspersed with different sequences of the other 14 tetragrams. Despite the strategic differences in exploring the maze, all organisms tested show that there is a single dominant strategy shared by others of that species. Regardless of the search pattern used, all species showed similar results in the ACF plots, with persistent, slowly decaying autocorrelation, indicative of long-lasting effect of choice on future choice selections. **(Fig. 3J-K_Supplementary_Time series analysis_Figure 4,5)**. These data collectively suggest that like zebrafish, mice and *Drosophila* did not search the test arena randomly, but in a systematic and deterministic way, demonstrating use of an underlying process of memory to recall previous turn choices, and guide subsequent turn patterns. This task provides further evidence of the suitability of the FMP Y-maze as a memory test for a range of model organisms **(Supplementary_Video. 2,3)**.

**Figure 4.**
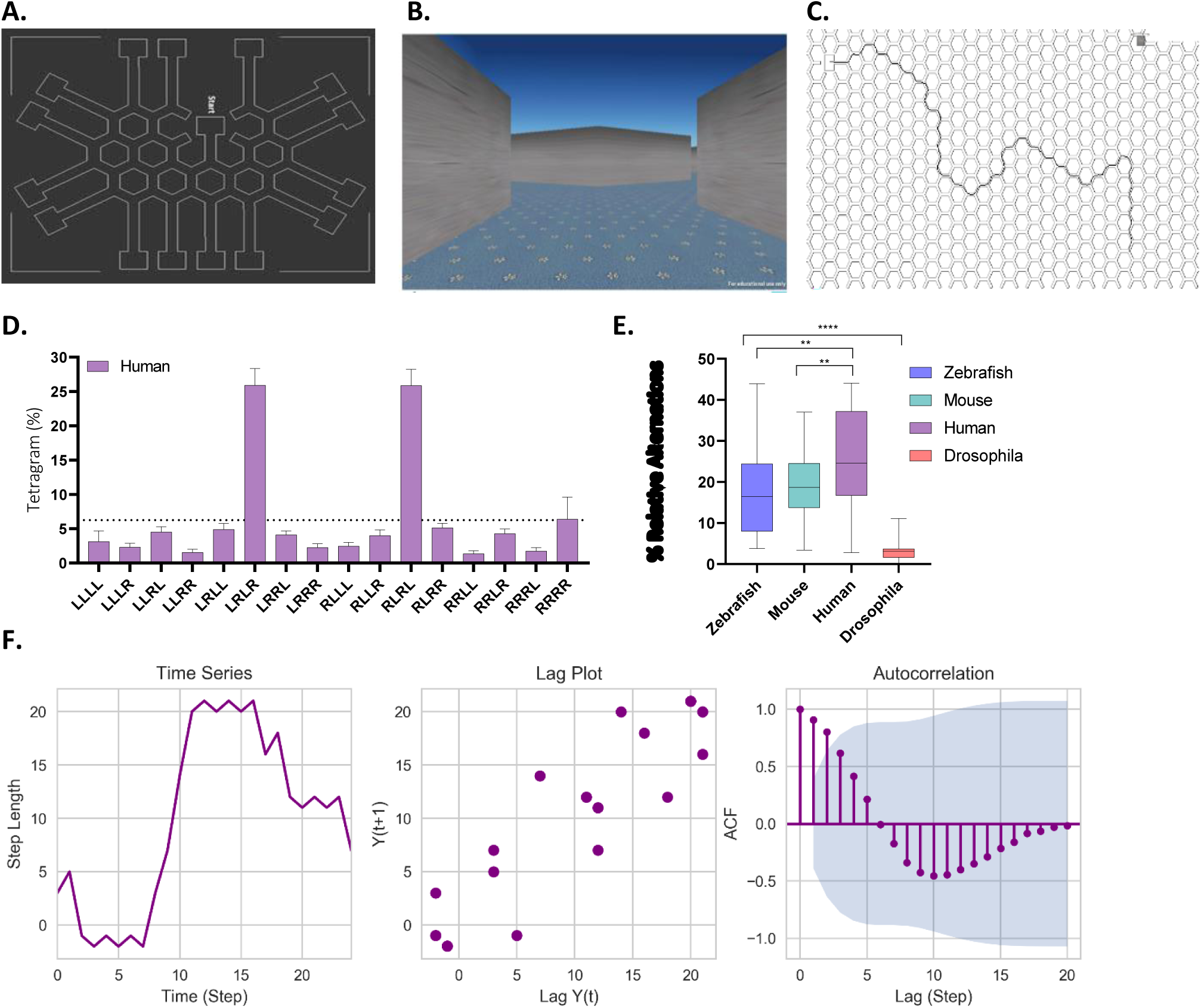
Virtual reality FMP Y-maze for humans. (**A**) Schematic of maze structure showing interconnected Y-shaped mazes, each of equal length and diameter. (**B**) In maze view from a participant’s perspective. (**C**) Line trace of a single participant based on the x,y coordinates from 5 minutes of exploration. (**D**) Tetragram frequency distribution of human participants from a 5 minute trial (n=24). (**E**) Relative means of alternations used in the FMP Y-maze of all organisms, demonstrating an increase in percentage use of alternation with complexity of organism, with *Drosophila* having the lowest use of alternations, increasing up to humans (**F**) Time series analysis of an individual participant showing time series plot, lag plot with weak positive correlation. ACF plot of the first 20 lags showing significant autocorrelation at lags 1 and 2 which exponentially decay to zero. Error bars represent ± SEM.

**Figure 5.**
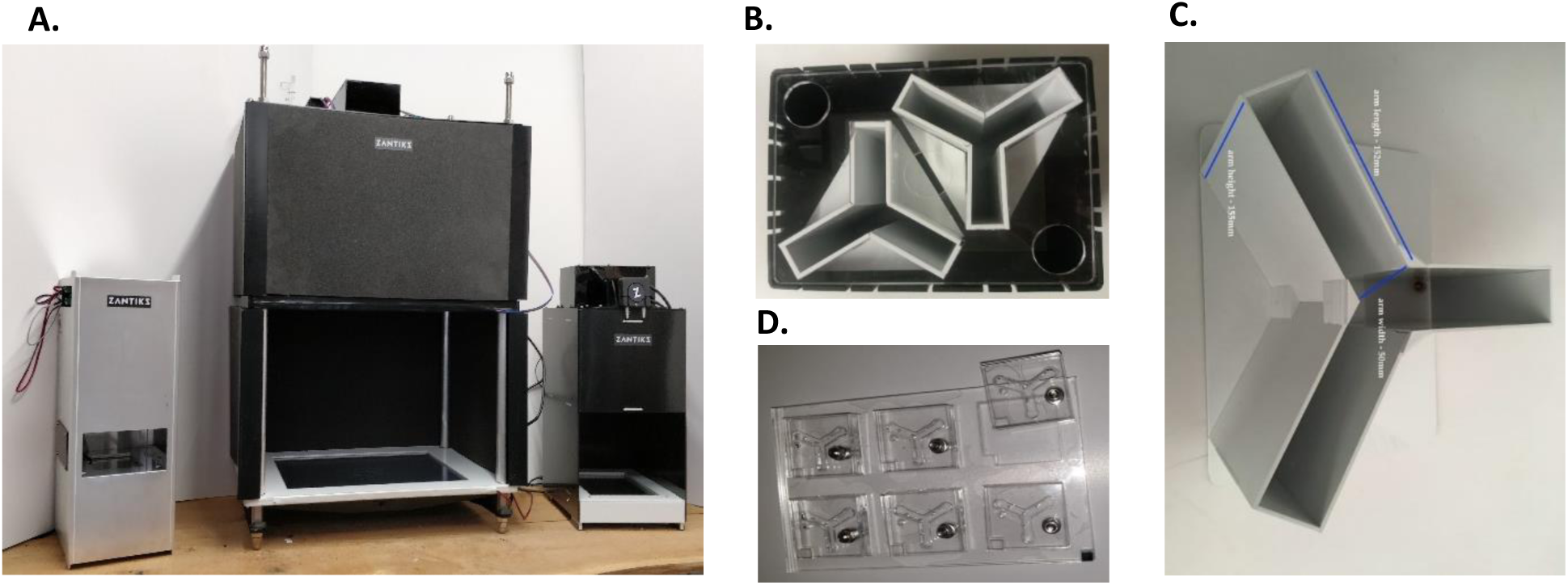
(**A**) Zantiks behaviour systems, from left to right, MWP system, LT system and AD system, used for *Drosophila*, mice and zebrafish respectively. Units are completely automated with a computer built into the base allowing for image/light projection and a camera positioned above, allowing live imaging of animals in the testing unit. This set up reduces experimenter disturbance during testing. (**B**) Zebrafish Y-maze inserts inside a black acrylic, water-tight tank. One fish per maze in 3L of aquarium water, equating to approximately 120mm water depth. (**C**) Mouse Y-maze insert. One mouse per maze. (**D**) *Drosophila* Y-maze insert, 6 identical mazes with sliding covers.

### Validating behavioural responses of humans to a virtual reality FMP Y-maze

In order to test the translational utility of the FMP Y-Maze to humans, we developed a virtual maze based on a honeycomb-layout **(Fig. 4A-B)**, thus requiring participants to navigate through a series of ‘Y’ shaped choice points. In order to make the test clinically relevant and useful for a variety of human testing conditions, we ran the task for a set time of 5 mins, at which point participants were automatically exited from the maze. In order to keep participants traversing the arena, they were instructed to find their way out before the time limit. Participants (male and female; age range 21-65) were recruited from staff and students at the University of Portsmouth. Participants could initiate the start of the trial when they were ready and use the arrow keys on a standard laptop keyboard to navigate around the maze **(Supplementary_Video.4)**. Interestingly, tetragram analysis revealed that humans used an almost identical strategy to mice and zebrafish, predominately comprising of alternations, which occupied ∼50% of the search strategy (one-way ANOVA; F(3, 164) = 60.88; P<0.0001) **(Fig. 4E)**. Despite the short run time, this prolific strategy was still detectable. On average, participants completed 39 steps (39 tetragrams) with a maximum of 68 and a minimum of 7 steps. The number of steps completed was substantially lower than any of the other animal models and was therefore based on 100 arm entries compared to 1000 arm entries for zebrafish, mice and *Drosophila* **(Fig. D_Supplementary_Time series analysis)**. Humans showed weak correlation in the lag plot and very few lags with significant autocorrelation in the ACF plot, lasting only one or two lags, then rapidly decaying to fluctuate around zero **(Fig. 4F)**. This indicates that the human FMP Y-maze exploration was characterised by choice selections that were only influenced by the immediate past. Based on the brevity of the trial and the limited number of turns this would be expected as the data set is not large enough to determine longer-term patterns. Our results have demonstrated the suitability of the FMP Y-maze as a test of memory, not just for animals, but also for humans, further supporting the theory of a common vertebrate strategy.

## Discussion

We demonstrate that the FMP Y-maze, when combined with tetragram analysis, is an effective tool for identifying search strategies and changes in memory function. There are several important benefits to using the FMP Y-maze. First, when used in conjunction with a human virtual system, it faithfully traverses cross-species behavioural phenotypes and has the ability to create a new dimension of memory assessment that could become bidirectional. Not only can behavioural changes in response to drug conditions be used to inform clinical trials, but specific behavioural traits of neuropsychiatric disorders in patients can be used to feed information back into animal models, a factor of great importance that is so often lacking in animal research^24^. Second, the task has the potential not only to reduce discrepancies between model organisms measuring the same conditions but could increase the success of translating treatments from animals to patients, a stumbling block that has seen a huge stall in the advancement of treatments of neuropsychiatric disorders in particular^25,26^. Third, the ability to detect impairment in cognition and memory at the most basic level, in the absence of training, habituation, reward bias or aversive conditions, creates a reliable test that can be run singly or as part of a battery of behavioural tasks assessing cognition and memory^8^. Fourth, the nature of the task is such that it can be applied by researchers from many backgrounds to measure aspects of memory without the need for in-depth knowledge of animal behaviour. Fifth, this task is non-invasive and low-impact on the animals, providing a task with a strong ‘3Rs’ (specifically refinement) justification. In summary, the FMP Y-maze lays the foundation of future translational research for a range of neurological disorders. This behavioural task could open new avenues of research into cognition and memory and allow cross-species comparisons with exceptional translational relevance.

## Methods

### Animals and housing

#### Zebrafish

A total of N=166 zebrafish (*Danio rerio)* of AB wild-type strain (4 months-old at time of testing) male and female (∼50:50) were bred in house and raised in the University of Portsmouth Fish Facility. Previous work from our lab has revealed no differences in search strategy between male and female zebrafish^27^. Fish were assigned at random to each treatment group from >10 groups of n = 11-18 fish per 2.8L tanks on a re-circulating system (Aquaneering Inc., San Diego, CA, USA). Sample sizes were calculated based on power analyses from effect sizes observed in pilot studies, and previous published work from our group ^28^. Room and tank temperatures were maintained at 25-27°C on a 14:10-hour light/dark cycle, water was aquarium treatment (dechlorinated) and pH is maintained at 8.4 (±0.4). Fish were fed on ZM fry food from 5 dpf until adulthood when they were moved onto a diet of flake food and live brine shrimp (ZM Systems, UK) 3 times/day (once/day at weekends). On completion of the behavioural testing fish were culled using Aqua-Sed anaesthetic treatment (Aqua-Sed™, Vetark, Winchester, UK) in accordance to manufacturer guidelines.

#### Mice

A total of N=16 C57BL/6 mice (*Mus musculus)* wild types (6-8 weeks old at the time of testing), male and female (50:50) were bred in house and raised in the University of Portsmouth Animal facility. Mice were housed in Allentown IVC racks and kept at 21°C (±2°C), 55% humidity (±10%) on a 12:12-hour light/dark cycle. Mice were fed on a diet of irradiated SDS RM3 pellets, with food and water available ad libitum. Following use, mice were retained as breeders in the University facility.

#### Drosophila

A total of N=30 Canton S wild-type (#64349) *Drosophila melanogaster* (6-7 days old at the time of testing), male and female (50:50), were obtained from Bloomington *Drosophila* Stock Centre, Indiana, USA. Flies were kept at 25°C with an average humidity of 60-80% on a 12:12-hour light/dark cycle. Flies were housed on ready mixed dried food (Phillip Harris, UK). Flies were collected via light CO_2_ anaesthesia and were allowed 48 hours recovery before behavioural testing was conducted. Following completion of the task *Drosophila* were culled using absolute ethanol.

### Ethical statement

All animal experiments were carried out following scrutiny by the University of Portsmouth Animal Welfare and Ethical Review Board (AWERB), and under a project licence from the UK Home Office (Animals (Scientific Procedures) Act, 1986) [PPL: P9D87106F]. For human experiments, ethical approval was granted by the University of Portsmouth Science Faculty Ethics Committee (SFEC), reference number SFEC 2019-062. Participants signed a consent form prior to participating, and only age and gender were recorded for each participant.

#### Pharmacological treatments

To examine the effects of MK801 (Sigma-Aldrich), scopolamine (Sigma-Aldrich), SCH-23390 (Tocris) and sulpiride (Sigma-Aldrich) on performance in the FMP Y-maze, fish were randomly allocated (from >10 groups of age-matched stocks in our fish facility) to one of our drug treatment groups with ∼13 fish assigned per group (n= 18 control, n= 13 0.1 mg/L MK801, n= 13 0.75 mg/L MK801, n= 13 2.0 mg/L MK801, n= 13. 0.25 mg/L scopolamine, n= 13 0.5 mg/L scopolamine, n= 13 1.0 mg/L scopolamine, n=12 0.5 mg/L SCH-23390, n=12 1.0 mg/L SCH-23390, n=12 mg/L 1.5 mg/L, n=12 5 mg/L sulpiride, n=11 10 mg/L sulpiride, n=11 20 mg/L sulpiride). Concentrations used were based on previously published works as well as range-finding pilot experiments in our laboratory ^5,29–32^. Fish were netted from home tanks and placed in 400 mL beakers containing 300 mL of either drug or fish water for 1 h. During pre-treatment, fish were isolated visually. This isolation avoided any impact of conspecifics or experimenters on treatment response. Following pre-treatment, fish were netted and placed in the FMP Y-maze (one fish per maze, 2 mazes per tank; **Supplementary_1.1_Figure. 1**) filled with 3L of fish water for 1 h. Data, comprising logs of entries into each arm, were automatically recorded throughout the 1 h period, minimising experimenter interaction. Following completion in the FMP Y-maze fish were immediately euthanized.

## Materials

Behavioural testing of all animals was conducted using a commercially available, fully integrated testing environment; the Zantiks MWP system for *Drosophila*, AD system for adult zebrafish, LT system for mice (Zantiks Ltd., Cambridge, UK, see **Fig. 5A**). Zebrafish were tested in a white acrylic Y-maze insert of two identical mazes (provided with the AD Zantiks base package) fitted into a black water-tight tank with a transparent base (https://www.zantiks.com/products/zantiks-ad) **(Fig. 5B)**. Mice were tested in a stand-alone white acrylic Y-maze insert with transparent base (provided with the LT Zantiks base package) (https://zantiks.com/products/zantiks-lt) **(Fig. 5C)**. *Drosophila* were tested in a clear acrylic Y-maze insert of 6 identical mazes. Each maze had a sliding cover with a hole which could be moved over the maze as an entry point for introducing the fly (extra for the MWP Zantiks unit) fitted into white opaque holding base for consistent maze alignment (https://zantiks.com/products/zantiks-mwp) **(Fig. 5D).** Mazes had equal arm length and angle. Maze dimensions were as follows; L50, W20, H140 (mm)-zebrafish, L152, W50, H155 (mm)-mice, L5, W3, H4 (mm)-*Drosophila*. Mazes were place into their respective Zantiks behaviour units, one maze for mice, two mazes for zebrafish and 6 mazes for *Drosophila*. Each system was fully controlled via a web enabled device. Filming was carried out from above, which allowed live monitoring within the behaviour system. Previous data from studies in the FMP Y-maze and continuous Y-maze indicated that extra-maze cues that allowed the fish to orientate did not affect exploration behaviour^28^. No cues were available in the mazes, and light levels were kept to a maximum of 2 lux during exploration, limiting the use of egocentric cues for navigation.

### Procedure

#### FMP Y-maze exploration task

The protocol was carried out as described in our previous papers^27,28,33^. Briefly, animal handling and experimenter visibility were both kept to a minimum. Fish were netted from home tanks, or from drug treatment beakers and placed directly in the maze. Mice were transported from home cage to maze using clear, plastic tubes that were kept in their home cages, preventing any direct handling of mice prior to the task. *Drosophila* were guided into a pipette tip and then tapped gently into the maze through a hole in the lid which could be moved over the maze to get flies in and on entry moved away from the maze to prevent escape. All animals were recorded in the maze in dim conditions (1-2 lux) for 1 h. As with our previous study, data were first output as a time series of arm entries and exits, normalised (proportions of total turns) and analysed according to 16 overlapping tetragrams (RLLR, LLRR, RRRL, etc.) of which particular note was taken with regard to search strategies termed alternations (RLRL, LRLR) and repetitions (LLLL, RRRR), having previously seen that these are most notably affected by different treatments. If the fish were adopting a random search strategy, it would be predicted that the distribution of tetragrams over a 1 h period would be approximately stochastic (i.e., the relative frequency of each tetragram ∼6.25%), and the data would generate autocorrelation plots equivalent to white noise (all lagged data points would fall below the 95% confidence interval).

#### Human Virtual FMP Y-maze

A honeycomb maze which presented multiple Y-shaped choice points formed the human virtual FMP Y-maze **(Fig. 4A-C)**. Participants were told that they could begin the trial when ready, and using the arrow keys on the keyboard of a laptop, try to find their way out of the maze before the time ran out. Participants were free to explore the maze for 5 minutes, after which they were automatically logged out. Turn directions were logged as x,y coordinates which were converted into left and right turns which could subsequently be transformed into tetragrams **(Supplementary_1.1)**. The human virtual FMP Y-maze was developed by Antony Wood, University of Southampton, using the 3DSMax 2012.

### Statistical Analysis

#### Tetragram analysis

Tetragrams were extracted from raw data via a custom-designed excel spreadsheet for analysis. First, to compare the difference between repetition and alternation across time, two-way RM ANOVA was used followed by Bonferroni-corrected pairwise comparisons. Second, to assess whether there were any effects of MK801, scopolamine, SCH-23390 or sulpiride on total activity (total number of turns in one hour), pure alternations (LRLR+RLRL), pure repetitions (RRRR+LLLL), right turns or left turns, we fitted generalized linear mixed effects model (Poisson distribution, log link), with concentration and time as fixed factors, and ID as a random effect (to account for non-independence of replicates). The results were considered significant when p ≤ 0.05. Descriptive statistics are calculated from raw data and expressed as means ± standard error of the mean (S.E.M) **(Supplementary_1.3_Tetragram analysis)**.

#### Time series analysis

Tetragram sequences were used to define step length and fix time intervals of the discrete time series. Each sequence was arbitrarily assigned a value ranging from 1 to 8. Left-dominant sequences were arbitrarily denoted as negative, whilst right-dominant sequences were positive **(Supplementary_Table. 1)**, from this point on referred to as ‘steps’. Each step was assumed equal time, therefore each observation in the time series was one tetragram sequence or the equivalent of one step. The analysis for zebrafish, mice and *Drosophila* were based on 1000 arm entries, sequentially divided into overlapping sequences of four arm entries, resulting in a total of 250 steps, *n*=250 time points. Human participants, due to the shortened trial time, completed significantly fewer arm entries; therefore, the total number of steps was *n=25*. Zebrafish treated with drugs, which variably affected locomotion, had a reduced average number of arm entries, therefore the total number of steps used for analysis was *n=150*. The limit was chosen arbitrarily for consistency only. Animals with more than 10 steps of missing data were excluded from subsequent time series analysis. Animals with fewer than 10 missing steps had zeros replacing missing values to make up the total number of steps required. The cumulative sum of steps were used to determine the relationship between successive observations and identify if steps were taken randomly and completely independent of one another. This was tested by computing the lag plot and autocorrelation function (ACF) using a custom designed script in MATLAB **(Supplementary_1.4_Time series analysis)**.

## ACKOWLEDGMENTS

We thank Antony Wood from the University of Southampton for allowing us use of the human virtual FMP Y-maze. This work has been supported by the University of Portsmouth Science Faculty Studentship, the Foundation for Liver Research, Alzheimer’s Research UK, Coordenação de Aperfeiçoamento de Pessoal de Nível Superior - Brazil (CAPES), University of Portsmouth and the Society for the Study of Addiction, University of East London.

## AUTHOR CONTRIBUTIONS

M.C., B.D.F., S.D.M., J.D.S. and M.O.P. designed the project; E.S.R. designed and developed the concept of the human virtual FMP Y-maze and extrapolated data; M.C., B.D.F. and D.C.R., performed the experiments and M.C., B.D.F., and M.O.P. analysed behavioural data; M.C. analysed time series data; and M.C. wrote the manuscript and all authors discussed the results and commented on the manuscript.

