## Supplemental methods for "A Test of Memory: The Fish, The Mouse, The Fly And The Human"

**Supplementary Information.**

**1.1) Adapted Y-mazes for different model organisms.**

**Figure 1.**

**A.** **B.** **C.**

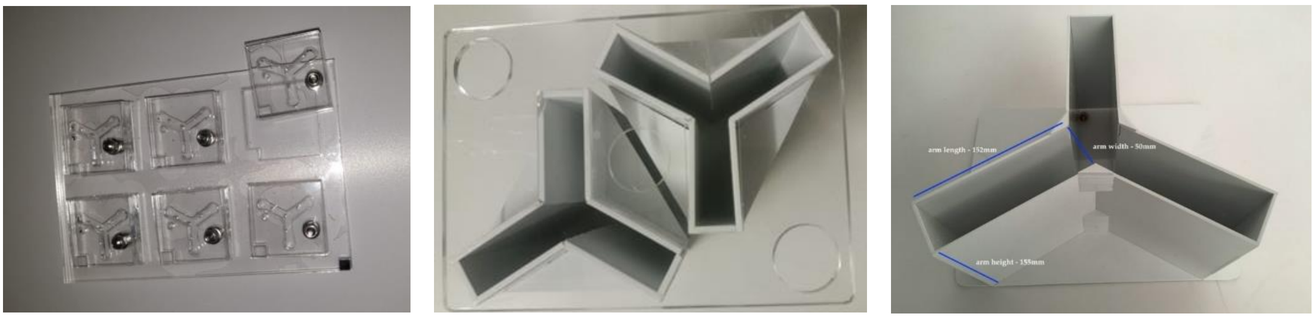

(**A**) Fly-maze with sliding cover to prevent escape. 6 Y-mazes can be run simultaneously. (**B**) Zebrafish maze, 2 Y-mazes can be run simultaneously, L50, W20, H140 (mm). Mazes are inserted into a water-tight tank which is filled with 3L of aquarium water. (**C**) Mouse Y-maze, only a single mouse can be run at a time, L152, W50, H155 (mm).

**Table 1.** Tetragram analysis was based on a series of 16 unique, overlapping sequences of left and/or right turns. Below is a list of each of the tetragrams used to conduct this analysis with reference to key strategies and the associated term.

| Sequence | Term | Step length | Sequence | Term | Step length |
|---|---|---|---|---|---|
| LLLL | Repetition | -8 | RLRL | Alternation | 1 |
| LLLR | | -7 | RLLR | | 2 |
| LLRL | | -6 | RRLL | | 3 |
| LRLL | | -5 | LRRR | | 4 |
| RLLL | | -4 | RLRR | | 5 |
| LLRR | | -3 | RRLR | | 6 |
| LRRL | | -2 | RRRL | | 7 |
| LRLR | Alternation | -1 | RRRR | Repetition | 8 |

**1.2) Heat maps and line trace of FMP Y-maze.** Heat maps and line traces of arm use were reported for selection of animals from each species chosen at random and run in the FMP Y-maze and tracked using Ethovision software to obtain graphics. Below are examples of heat maps and line traces from each species.

**Figure. 2**

Zebrafish

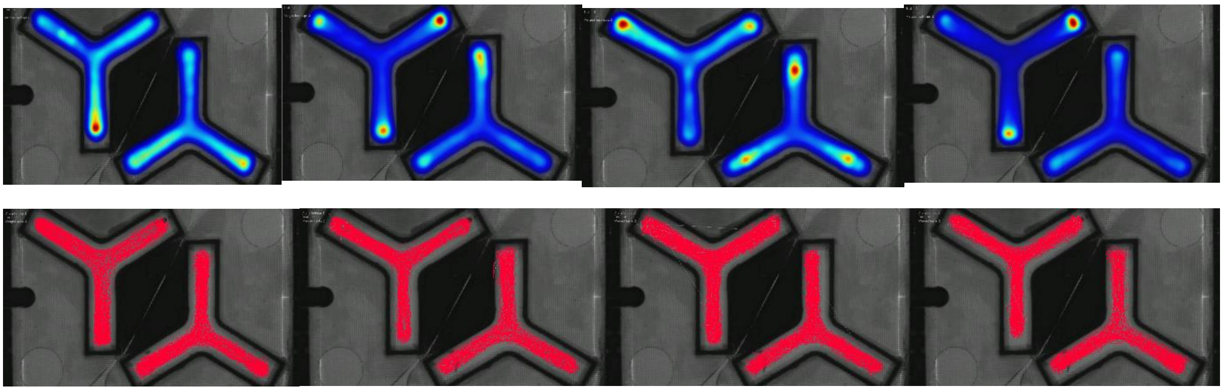

Mouse

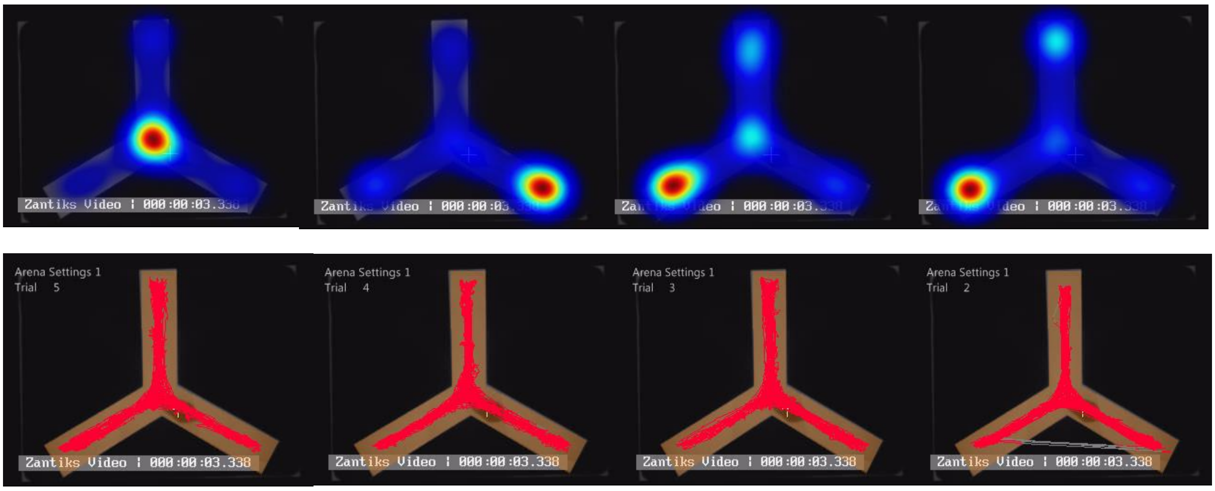


*Drosophila*

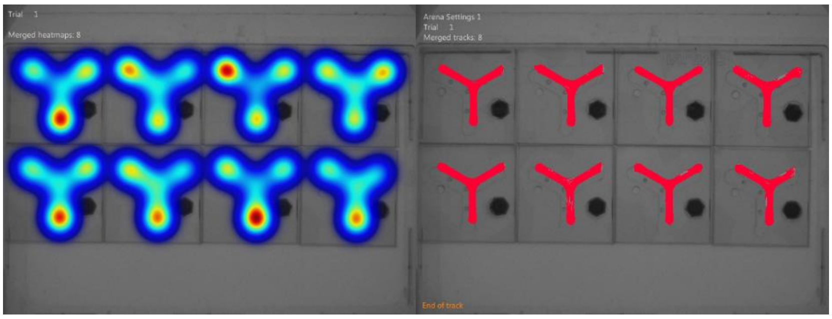

**1.3) Tetragram Analysis.** In a test paradigm consisting of two equally likely choice variants, left (L) or right (R) turn, we assume choice selection to be completely random. However, we know from behaviour of humans in guessing tasks [34,35], or animals in choice behaviour tasks, such as rodents in a T-maze preferring to alternate L and R turns despite equal arm reinforcement [4,36], that choices are never completely random. In a Markov process, a process of completely random events, the probability of choosing L or R depends only on the most recent choice [37]. For example, the probability of turning L would be:

$$P(L) = ½,$$

regardless of whether the previous turn had been L or R. Despite the overall process being random, it is possible to detect patterns in large data series by dividing sequences into groups of like-terms and using information theory to detect any departures from randomness[38]. Let $p_i$ be the probability of event *i* in a time series, such as the probability of turning L or R. Using general information theory formula the first order 'uncertainty' of turning L after previously turning R could be measured using:

$$L = \sum p_i \log_2 p_i,$$

where base 2 for the logarithm stipulates that from two equally-likely events (L or R), one choice (one unit of information) is transmitted to resolve the uncertainty of the occurrence of either choice. Relative uncertainty, $L_{max}$, is the ratio of observed L turns to maximum L turns, for the given number of alternatives, the compliment of this is:

$$1\text{-}L/L_{max}$$

Different levels of complexity can be used to determine the probability of turning L based on two previous turns, LR (digram of events), three previous turns, LRL (trigram), four previous turns, LRLR (tetragram), etc. The larger the number of alternative choices the greater the computational power required. Previous work has demonstrated that in human guessing tasks, examination of past events exceeding four or five choices becomes irrelevant when calculating the probability of a current event[38,39]. Therefore, in line with previous two-choice guessing task protocols, we have selected to concentrate on the use of tetragram sequences, limiting the number of alternatives to $2^4 = 16$ possible tetragram sequences. The information measure for a sequence of four turn choices for turning L is:

$$L_4 = L(\text{tetragram}) – L(\text{trigram})$$

**1.4) Time Series Analysis.** Time series, $\chi_n = \chi_1, \chi_2 \ldots\ldots\ldots\ldots X_k$ were defined as step length, ω(*k*), at discrete time, *k*, where *k* was representative of equal length time points comprised of tetragram sequences. Put simply, each point in the time series was equal to one tetragram, described as one step. Each experiment was made up of *n* time points. The autocorrelation lag coefficients of steps were calculated for each individual using step length, ω(*k*). ACF was computed in PYTHON using MATLAB[40]. The lag-1 autocorrelation for the corresponding time lag *k* is:

$$\text{ACF}(k) = \frac{\sum_{s-1}^{T-k}(\omega(s)-\bar{\omega})(\omega(s+k)-\bar{\omega})}{\sum_{s=1}^{T}(\omega(s)-\bar{\omega})^2},$$

where $\bar{\omega}$ is the mean step length for that individuals time series, ω(*k*). As the model demonstrated non-stationary and non-random properties, the usual calculation of confidence interval, $\bar{\omega} \pm 2\sigma \,/\, \sqrt{n}$ , where σ is the standard deviation, was not used. Instead the 95% confidence interval was based on a moving average calculated using the Bartlett test:

$$\frac{T = (n-k)\, ln\, \sigma_p^2 - \sum_{i=1}^{k}(n_i - 1)\, ln\, \sigma_i^2}{1 + \left(1, \left(3(k-1)\right)\right)\left(\left(\sum_{i=1}^{k} 1 \,/\, (n_i - 1)\right) - 1 \,/\, (n-k)\right)}$$

Where $\sigma_i^2$ is the variance of the *i*th group, *n* is the total number of steps, $n_i$ is the step length of the *i*th group, *k* is the number of groups and $\sigma_p^2$ is the weighted mean of the group variances, defined as:

$$\sigma_p^2 = \sum_{i=1}^{k}(n_i - 1)\sigma_i^2 \,/\, (n-k)$$

**Figure 3.** Individual zebrafish ACF plot of lag-1 coefficients showing 20 of 250 lags.

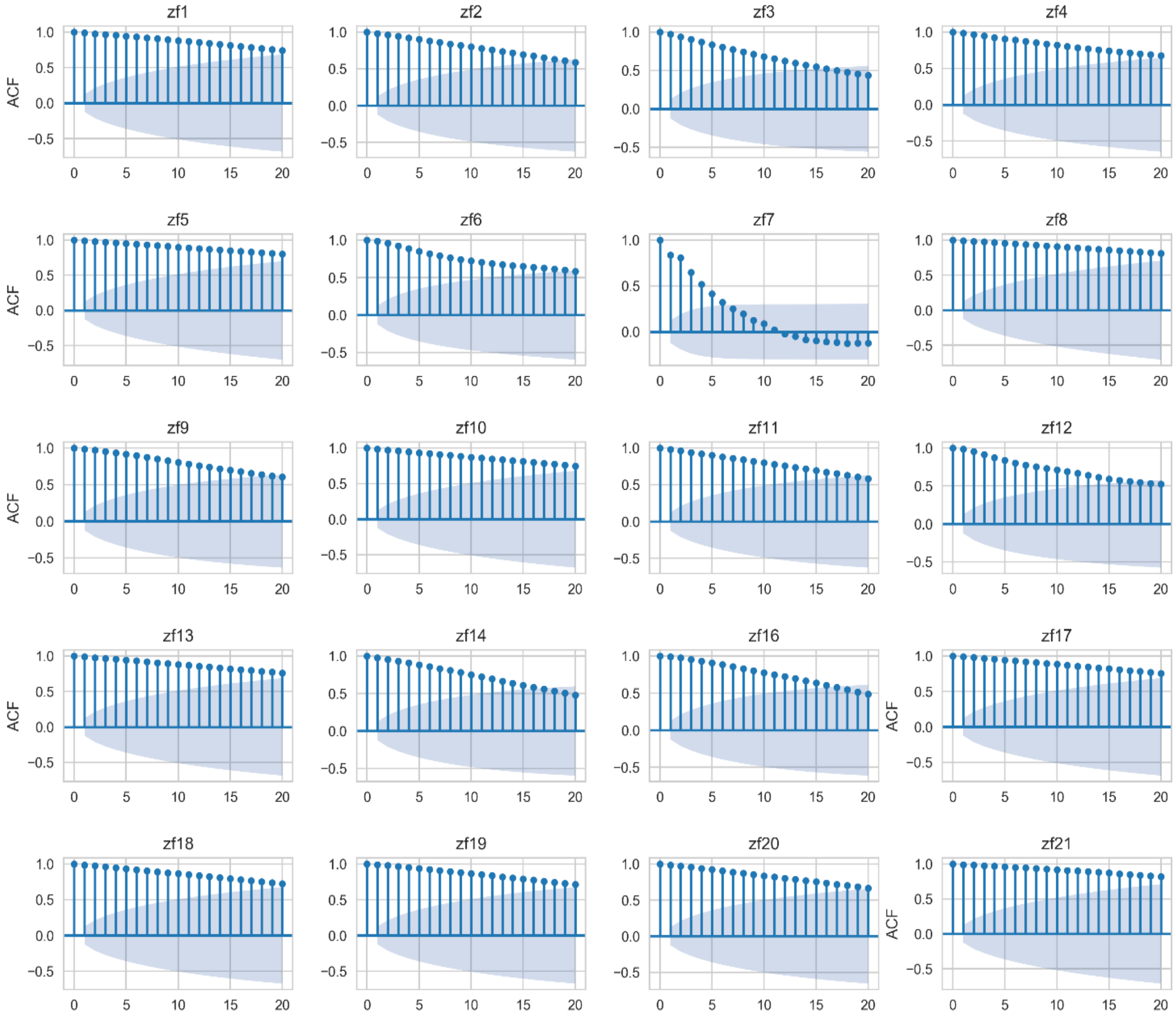

**Figure 4.** Individual mouse ACF plot of lag-1 coefficients showing 20 of 250 lags

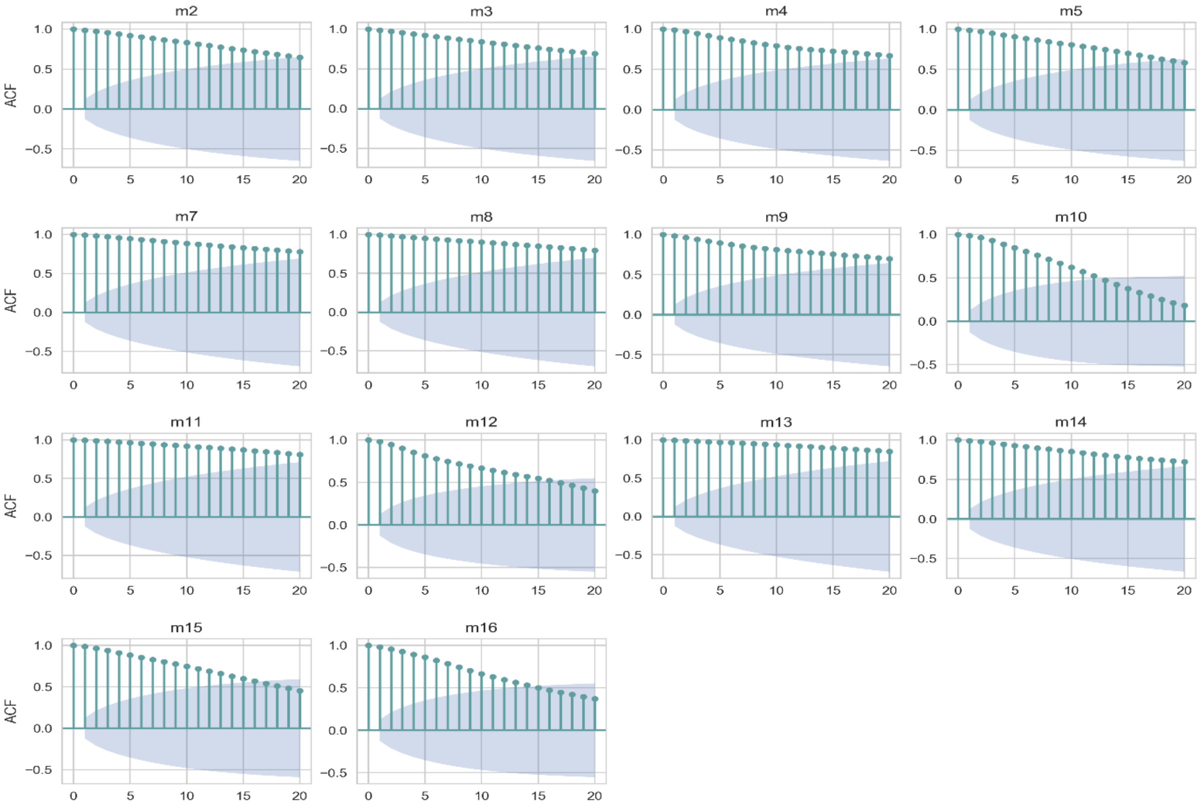

**Figure 5.** Individual *Drosophila* ACF plot of lag-1 coefficients showing 20 of 250 lags

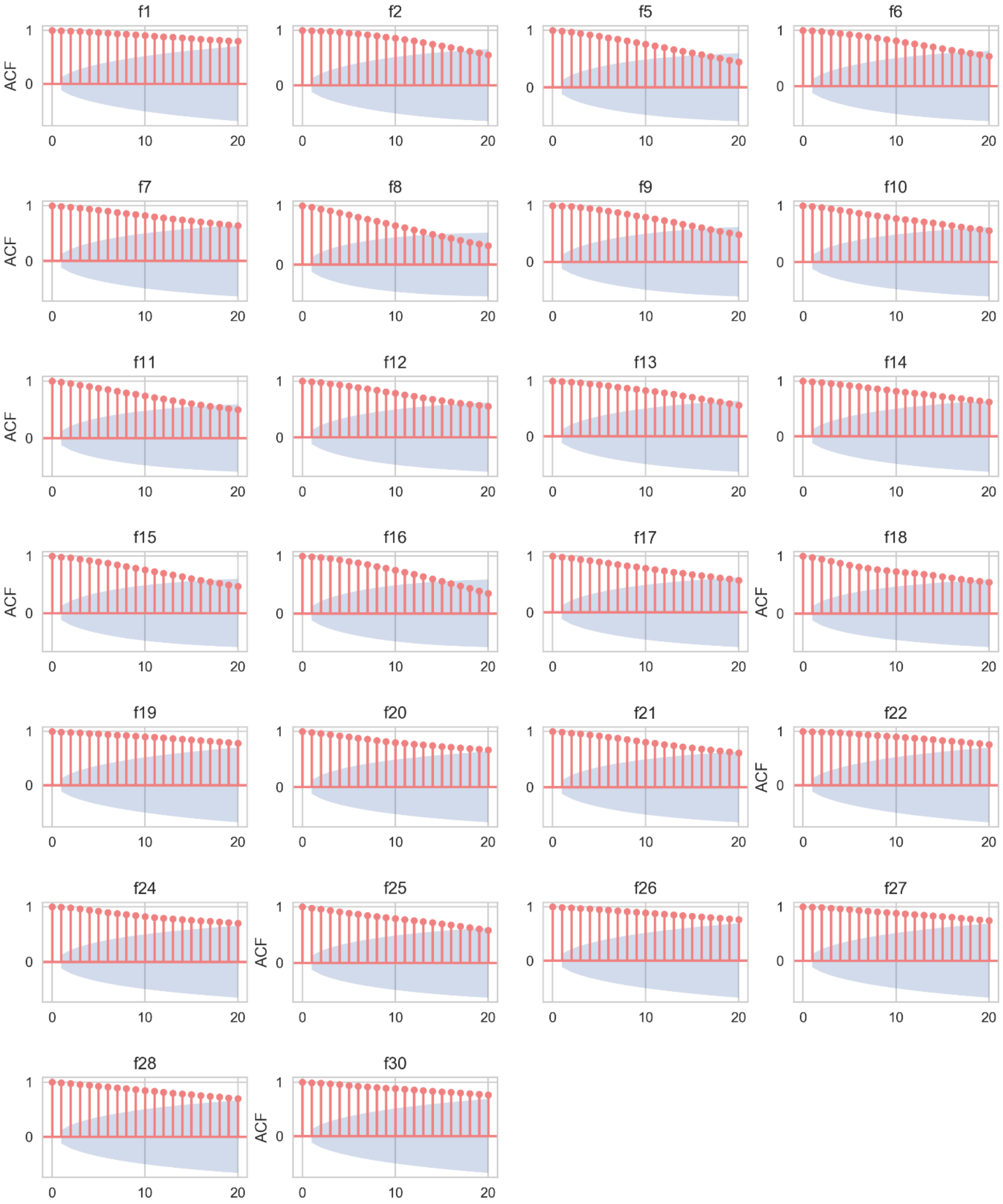

**Figure 6.** Individual human participant ACF plots of lag-1 coefficients showing 20 of 25 lags.

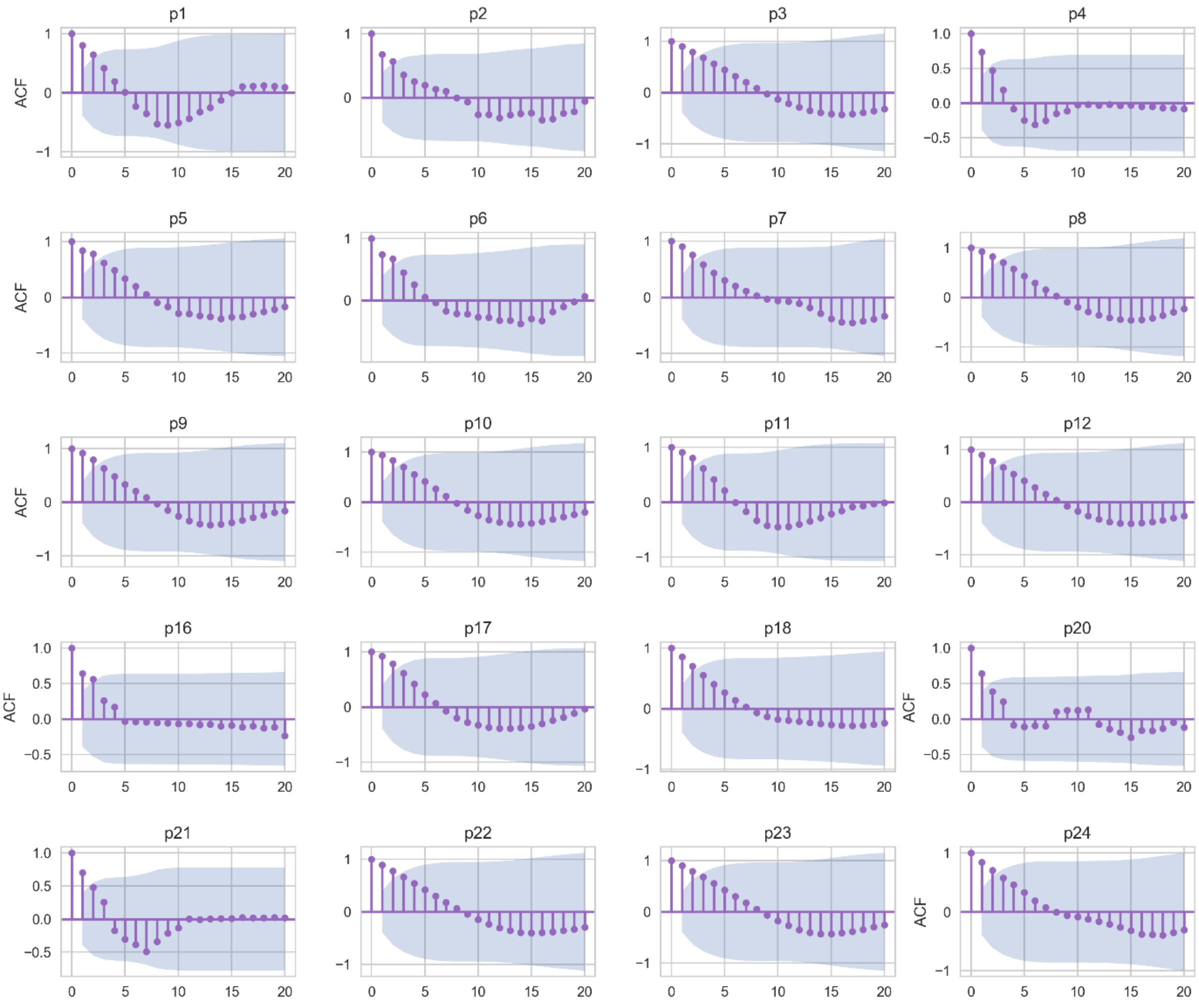

**Figure 7.** Individual zebrafish ACF plots of lag-1 coefficients showing 20 of 150 lags pre-treated with D1 antagonist SCH-23390 at (A) 0.5 mg/L, (B) 1.0 mg/L and (C) 1.5 mg/L

A.

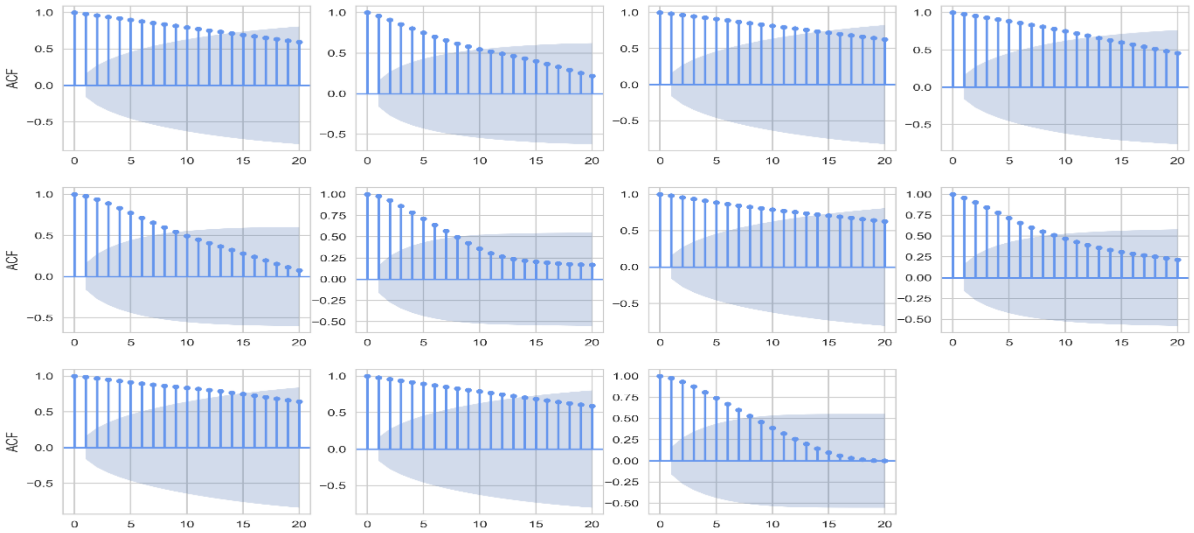


B.

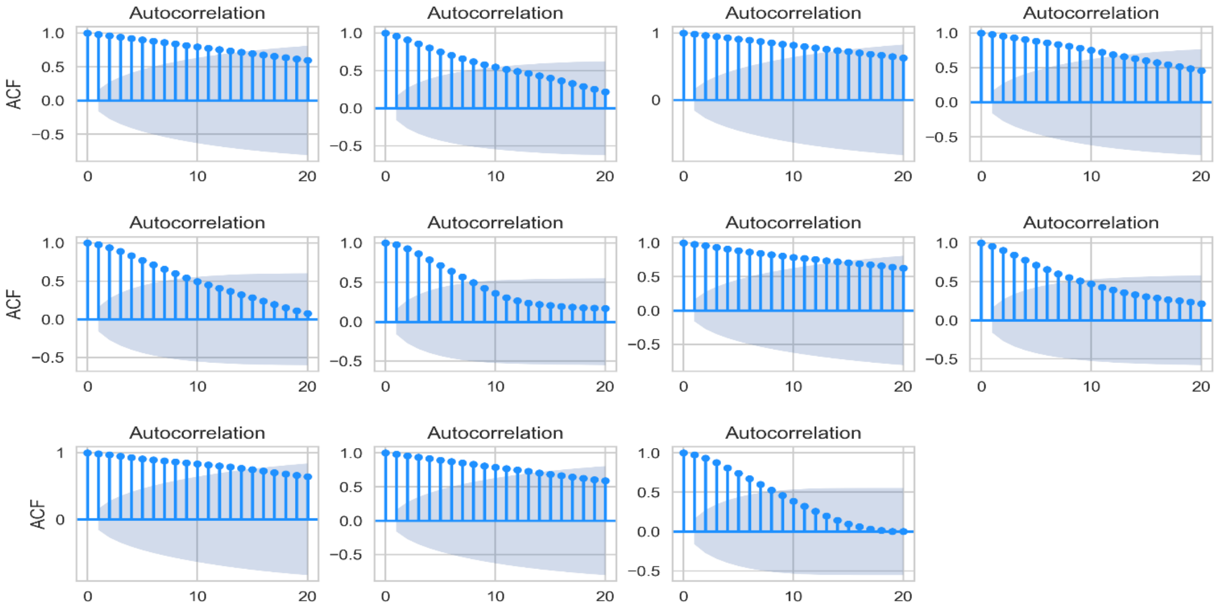


C.

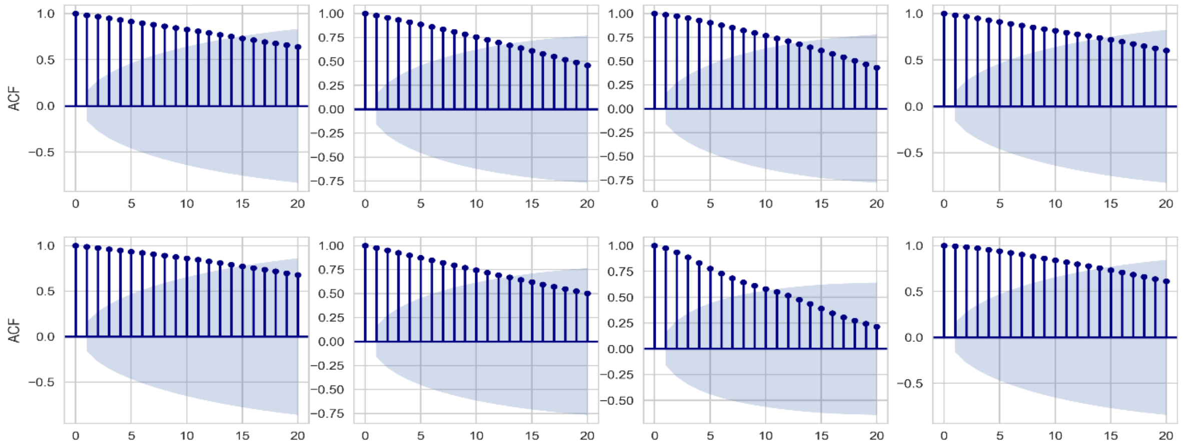

**Figure 8.** Individual zebrafish ACF plots of lag-1 coefficients showing 20 of 150 lags pre-treated with D2 antagonist sulpiride at (A) 5.0 mg/L, (B) 10.0 mg/L and (C) 20.0 mg/L.

A.

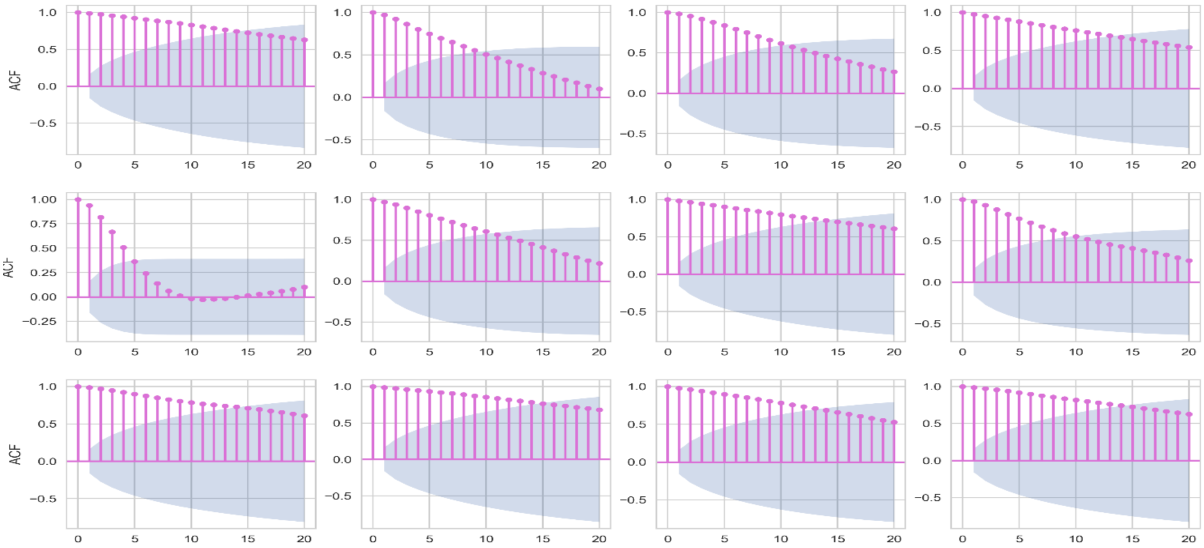


B.

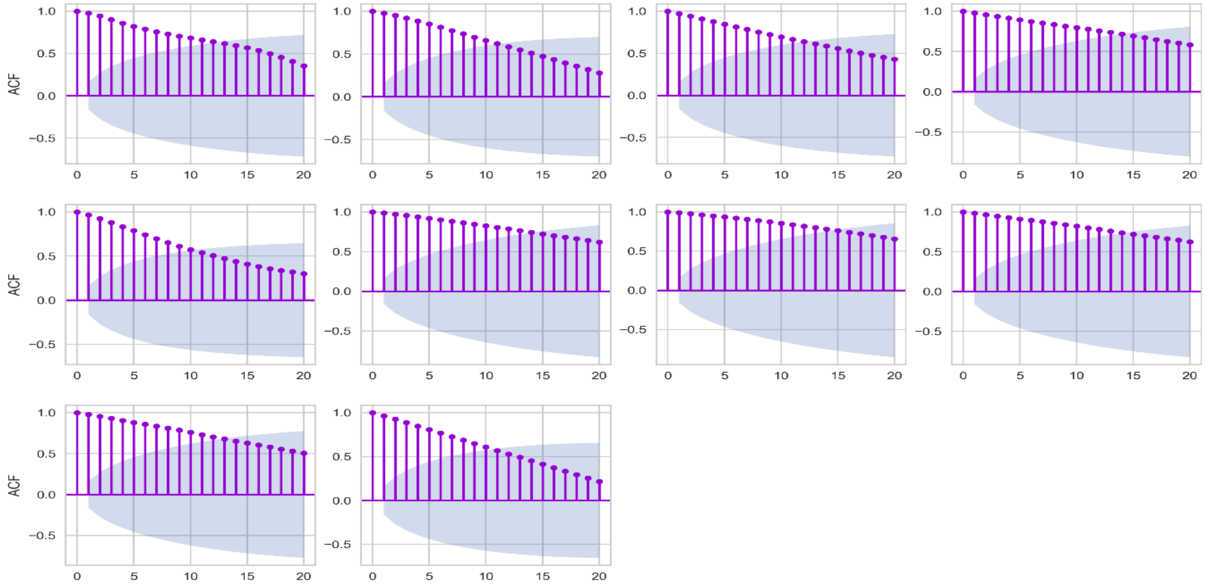


C.

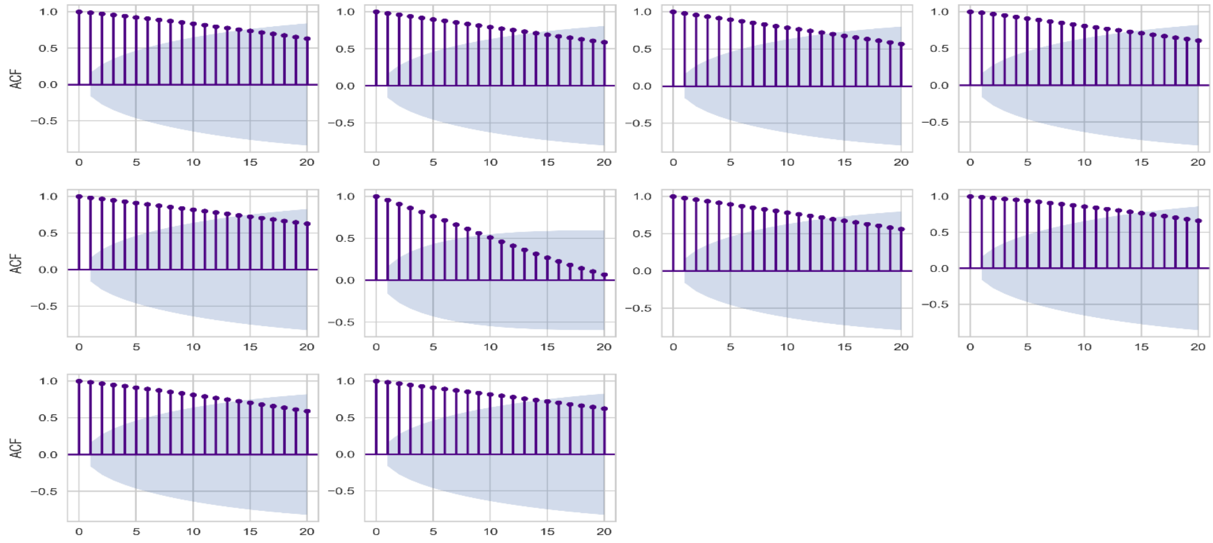

**Figure 9.** Individual zebrafish ACF plots of lag-1 coefficients showing 20 of 150 lags pre-treated with NMDA-r antagonist MK801 at (A) 0.1 mg/L, (B) 0.75 mg/L and (C) 2.0 mg/L.

A.

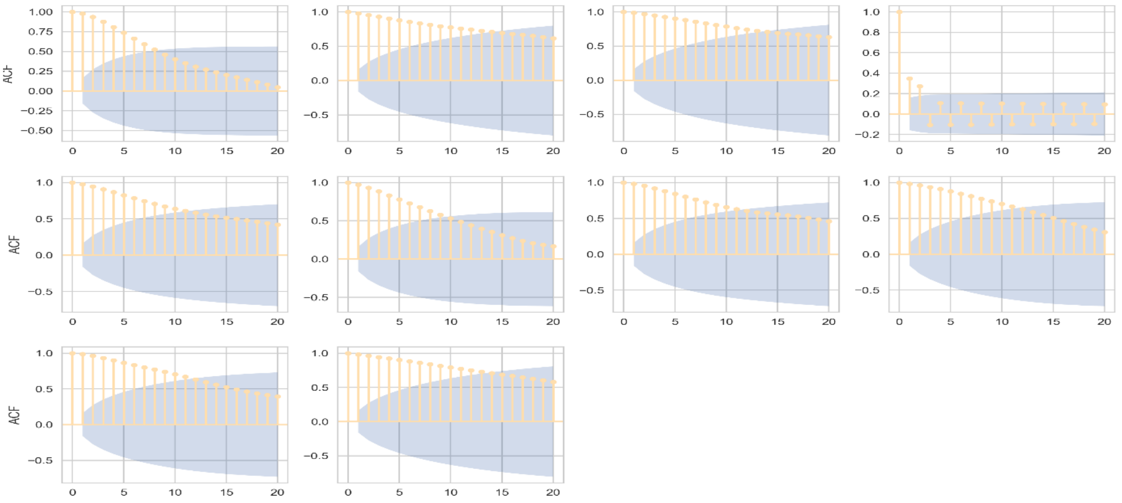


B.

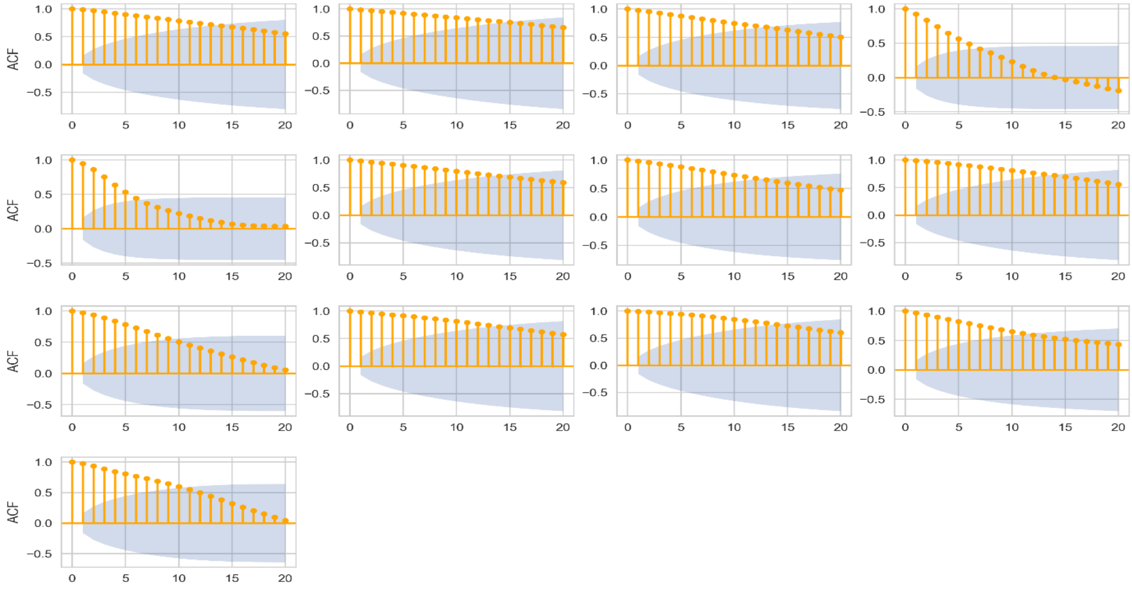


C.

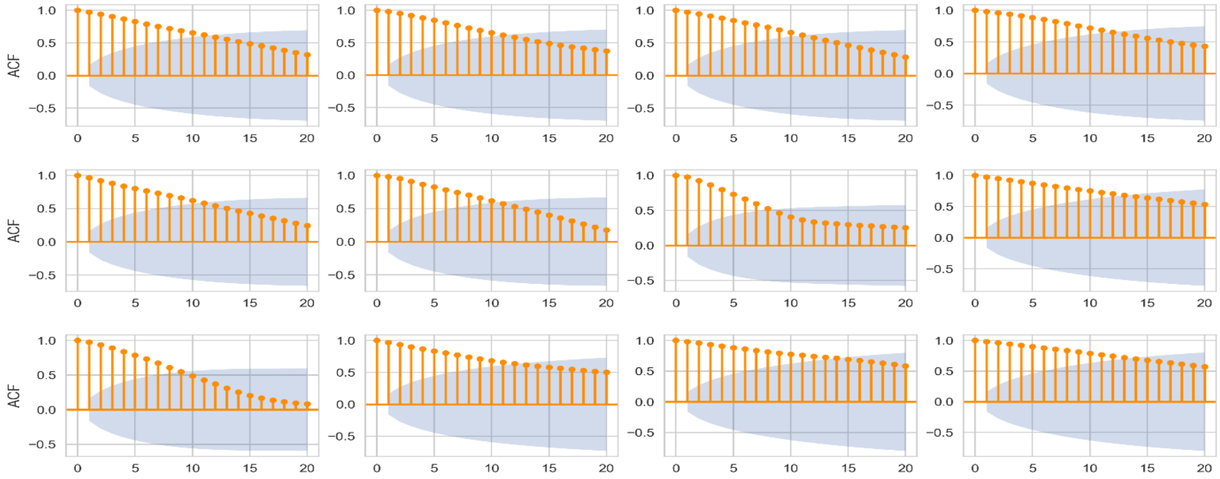

**Figure 10.** Individual zebrafish ACF plots of lag-1 coefficients showing 20 of 150 lags pre-treated with muscarinic-r antagonist scopolamine at (A) 0.25 mg/L, (B) 0.5 mg/L and (C) 1.0 mg/L.

A.

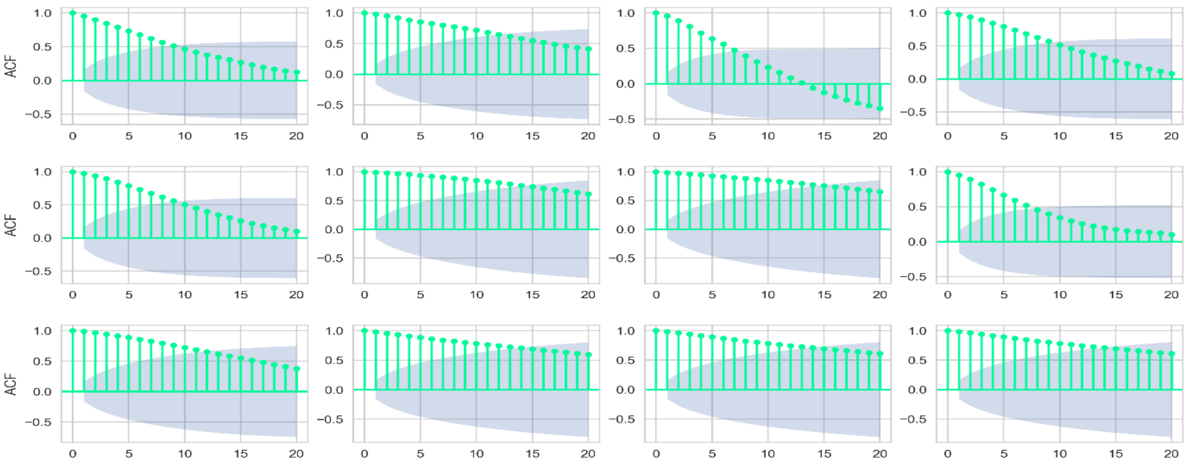


B.

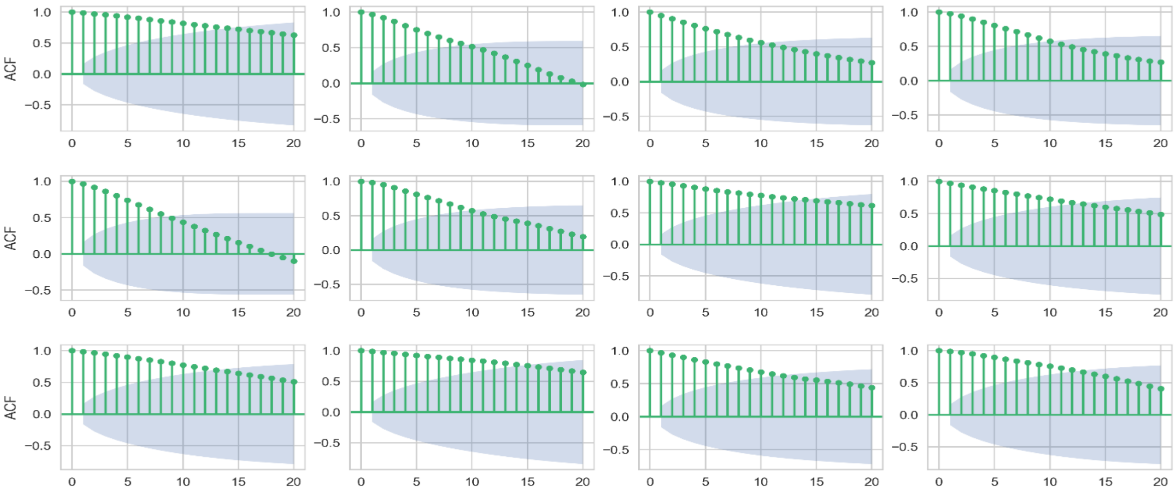


C.

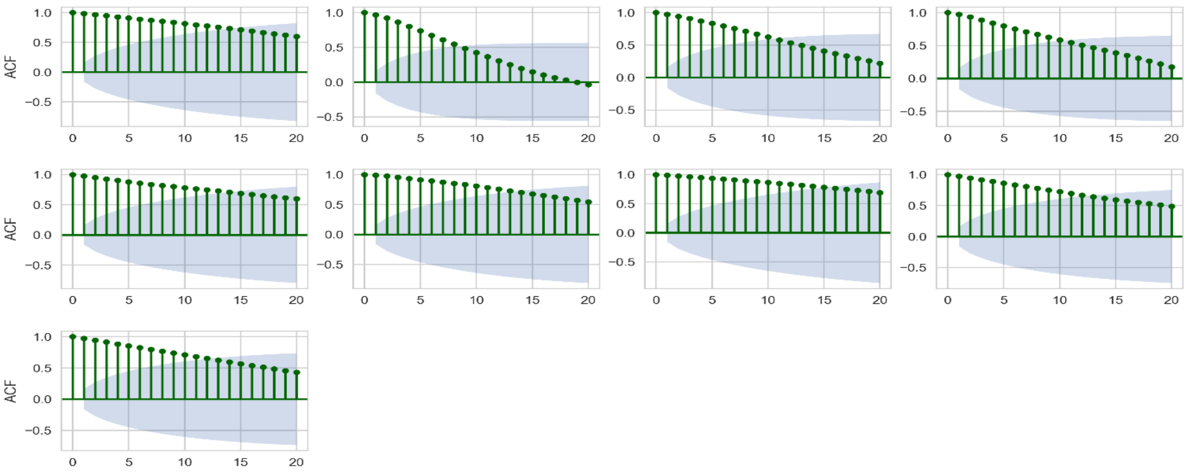

**1.5) Human exploration in the VR FMP Y-maze.** The human exploratory task presented in the virtual reality FMP Y-maze had superficial differences from the general model of exploration used in the animal model of the FMP Y-maze. The main differences were the maze design and the total exploration time. To prevent participants getting bored or ending the task prematurely, a complex of multiple Y-shaped mazes were combined to form a honey combe shaped maze allowing exploration of locations relative to their current position in the maze. Exploration time was limited to 5 minutes to prevent the task from becoming too time consuming and off-putting to participants. The brevity of the task easily lends itself to testing multiple participants in a short time.

**Video 1**. Zebrafish FMP Y-maze

**Video 1.1** Zebrafish FMP Y-maze – example of repetition and alternation strategies

**Video 2**. Mouse FMP Y-maze

**Video 3**. *Drosophila* FMP Y-maze

**Video 4**. Human VR-FMP Y-maze
